# Tackling emerging artemisinin resistance by modulating the defensive oxido-reductive mechanism of human malaria parasite by repurposing Nitrofurantoin

**DOI:** 10.1101/2023.04.18.537303

**Authors:** Sadat Shafi, Sonal Gupta, Ravi Jain, Rumaisha Shoaib, Akshay Munjal, Preeti Maurya, Abul Kalam Najmi, Shailja Singh

## Abstract

Oxidative stress mediated cell death has remained the prime parasiticidal mechanism of front line anti-malarial, artemisinin (ART). The emergence of resistant *Plasmodium* parasites characterized by oxidative stress management due to impaired activation of ART as well as enhanced ROS detoxification has decreased its clinical efficacy. This gap can be filled by development of alternative chemotherapeutic agents to combat resistance defense mechanism. Interestingly, repositioning of clinically approved drugs presents an emerging approach for expediting anti-malarial drug development and resistance management. Herein, we evaluated the anti-malarial potential of Nitrofurantoin (NTF), a clinically used antibacterial drug, against intra-erythrocytic stages of ART-sensitive (*Pf*3D7) and resistant (*Pf*Kelch13^R539T^) strains of *Plasmodium falciparum* (*Pf*), alone and in combination with ART. NTF exhibited growth inhibitory effect at sub micro molar concentration by arresting parasite growth at trophozoite stage. It also inhibited the survival of resistant parasites as revealed by ring survival assay. Concomitantly, *in vitro* combination assay revealed synergistic association of NTF with ART. NTF was found to enhance the reactive oxygen and nitrogen species as well as induced mitochondrial membrane depolarization in parasite. Furthermore, we found that exposure of parasites to NTF disrupted their redox balance by impeding *Pf* Glutathione Reductase activity, which manifests in enhanced oxidative stress, inducing parasite death. *In vivo* administration of NTF, alone and in combination with ART in *P. berghe*i ANKA infected mice blocked parasite multiplication and enhanced mean survival time. Overall, our results indicate NTF as a promising repurposable drug with therapeutic potential against drug sensitive as well as resistant parasites.

## 1. Introduction

Malaria, a protozoan infectious disease caused by *Plasmodium* species, is competent of inducing mild to serious infections in humans. Despite considerable therapeutic improvements, malaria still remains a global health burden due to emergence of drug resistant strains [1]. Artemisinin (ART)-based combination therapies (ACTs) are considered as the front line therapeutic interventions against malaria [2]. Importantly, the parasiticidal mechanism of ART have been linked to iron mediated free radical generation, thereby increasing oxidative stress on the parasite [3]. The accertion of intracellular oxidative stress induces oxidative damage to bio-molecules including lipids, nucleic acids and proteins thereby promoting cell death [4]. However, the global rise of ART-resistant *Plasmodium* strains, primarily in Eastern and Central India, South America (Fr Guiana), and East Africa [5], challenges this treatment. The resistance to ART relies on the parasite’s potential to attain a quiescent phase, with a significantly reduced metabolism [6]. Interestingly, the mechanism of ART resistance is linked primarily to *Pfk13* gene polymorphism [7], which incorporates several molecular changes in the resistant parasite [3]. The *Pfk13* gene mutation impairs the acquisition and breakdown of hemoglobin in ART resistant parasites [8], resulting in reduced heme-mediated ART activation in these parasites and so, a decrease in oxidative burden [9]. Additionally, resistant parasites demonstrate superior ability to maintain oxidative onslaught [10], leading to decreased efficacy to ART and jeopardizing the effect of partner drugs. This intensifies the need for developing novel lead molecules in the anti-malarial drug pipeline. Interestingly, drug repositioning presents an attractive and a viable fast-track drug development. Considering the emergence of ART resistance, it is imperative that compounds from existing therapeutic pool are scientifically screened to demonstrate their anti-malarial potential, and also identify partner candidates for ART based combinatorial regimes. Scientific reports have described nitro-aromatic antibiotics with promising anti-malarial efficacy against drug sensitive and resistant parasites [11,12]. Encouraged by the documented results, we investigated the chemotherapeutic potential of a conventional nitroaromatic antibiotic, Nitrofurantoin (NTF) for malaria, and further linked its anti-malarial efficacy as *Pf* Glutathione Reductase (*Pf*GR) inhibitor.

NTF is a drug of choice for treating acute lower urinary tract infections (UTI), chronic UTI, and for the suppression of catheter-associated bacteria [13]. It triggers bacterial cell death through oxidative stress induction, with effectiveness against both gram-negative and positive bacteria [14,15]. Notably, NADPH-dependent reduction of NTF generates nitroaromatic anion radicals, whose auto-oxidation generates free radicals, inducing oxidative stress on bacteria. Subsequently, this drug presents an excellent safety profile and managed to avoid resistance problems, despite several decades of use. This study reports the re-profiling of anti-bacterial drug, NTF against sensitive and resistant (*Pf*3D7 and *Pf*Kelch13^R539T^) strains. NTF exhibited a potent anti-plasmodial activity, alone as well as in combination with ART, against *Pf* both in *in vitro* and *in vivo*. Moreover, NTF treatment significantly impeded the survival of ring-stage parasites of resistant strain. Mechanistically, the potent growth inhibitory effect of NTF can be linked to oxidative stress induction by auto-oxidation as well as dysregulation of redox homeostasis.

## 2. Material and Methods

### 2.1 In vitro Parasite culture

*Pf*3D7 and *Pf*Kelch13^R539T^ strains were cultured at 2% hematocrit in O+ erythrocytes (Rotary Blood Bank) in RPMI 1640 medium (Invitrogen, USA), supplemented with Albumax I (0.5%), Hypoxanthine (50 mgL^-1^), Sodium bicarbonate (2g/l) and Gentamicin (10 mg/L), in a 37°C mix gas setting (90% N_2,_ 5% CO_2_ and 5% O_2_) [16]. Parasite culture was monitored regularly by Giemsa (Sigma, USA) staining and synchronized with 5% sorbitol. Late staged parasites were further purified by using 65% percoll.

### 2.2 Growth inhibition assay

To assess the *in vitro* growth inhibitory potential of NTF, synchronised ring stage parasites (*Pf3D7* and *PfKelch*13^R539T^) at 0.8-1% parasitemia were incubated with NTF at concentrations ranging from 1.25μM-40 μM. ART served as a standard drug, while untreated culture was used as a negative control. Following 66h incubation at 37ºC, parasitemia was assessed by counting 2000 cells in giemsa stained blood smears of each assay condition. The growth inhibition was computed by the following equation:

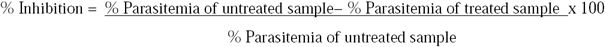

### 2.3 Stage specific growth inhibition assay

The stage-specific growth inhibitory potential of NTF was assessed in accordance with a standard protocol [17]. In brief, synchronized *Pf*3D7 culture (0.8-1% parasitemia) was exposed for 6h to varying concentrations of NTF at ring stage (0 to 6 h), trophozoite stage (28 to 34 h), and schizont stage (42 to 48 h). Following 6h incubation, test compound was removed by washing the parasites with complete medium, and then further cultured till trophozoite stage of next cycle. Untreated culture served as control. Percentage chemo-suppression was estimated by comparing parasitemia level of drug-treated with untreated control.

### 2.4 Detection of Intracellular reactive oxygen species by DCFDA

Synchronized trophozoites (8-10% parasitemia) were incubated for 30 minutes with 20 μM Dichlorodihydrofluorescein diacetate (DCFDA) [18], followed by treatment with NTF and ART at 4xIC_50_ for 6 h. Untreated parasites served as negative control. Following washing, the cells were mounted onto a glass slide and visualized under fluorescence microscope (Olympus). The mean fluorescence intensity (MFI) was estimated for 15 cells each from treated and untreated samples, using NIS elements software and plotted.

### 2.5 Estimation of Nitric Oxide Level

The effect of NTF on production of reactive nitrogen species (RNS), estimated as nitrite (NO2−) level was determined using Griess reagent kit (Invitrogen). Trophozoites at 8-10% parasitemia were exposed to varying concentrations of NTF for 24 h. Following incubation, culture supernatant from each sample was mixed with Milli-Q water and Griess reagent (N-(1-naphthyl)-ethylenediamine: sulfanilic acid) and incubated in 96 well plate at RT for 30 minutes. Deionized water was used to prepare a blank sample. The absorbance of each sample at 548 nm was quantified using a microplate reader and plotted.

### 2.6 *Detection of Mitochondrial Membrane Potential* (∆Ψ_m_) *using JC-1 dye*

The ∆Ψ_m_,, an indicator of cell death, in parasites was detected using JC-1 dye, in accordance with the previously described protocol [19]. Briefly, trophozoites at 8-10% parasitemia were incubated with NTF and ART (positive control) at 4xIC_50_ concentration for 6 h. After incubation, parasites were washed with PBS, and then incubated for 30 minutes with JC-1 dye (5 μM) in dark. Following washing, cells were mounted onto a glass slide and visualized under fluorescence microscope. MFI was assessed of each channel using NIS elements software.

### 2.7 Interaction study of PfGlutathione Reductase with NTF

#### 2.7.1 Architecture of Pf Glutathione Reductase

Amino acid residues of *Pf*Glutathione Reductase (*Pf*GR) involved in the interaction with FAD, NADPH and GSSG, constituting the respective catalytically active sites/pockets; and, those involved in forming a cavity/pocket at the dimer interface, were employed to illustrate overall architecture of the protein [20–22]. This was done by using Illustrator for Biological Sequences (IBS 1.0.3) [23]. X-Ray diffraction based structural model of *Pf*GR (PDB ID: 1ONF; available at resolution: 2.6 Å) [24], was procured from the Protein Data Bank (PDB).

#### 2.7.2 In silico binding analysis of Nitrofurantoin with PfGR

Structural Data Format (SDF) file of NTF was procured from PubChem. Energy-minimized 3D structural model of NTF was produced by utilizing Chem3D Pro 12.0 (https://perkinelmerinformatics.com/products/research/chemdraw/). *In silico* interaction analysis was done via Autodock Vina Tools 1.5.6 [25], as reported previously by our group [26,27,p.1], to rationalize the inhibitory activity of NTF against *Pf*GR. The plasmodial protein harbors amino acid residues involved in the interaction with FAD, NADPH and GSSG, constituting the respective catalytically active sites; and, those involved in forming a pocket at intersubunit-contact [20–22,28]. Both cavities of *PfGR* have earlier been reported to bind numerous non-competitive inhibitors, and are targets for selective drug design [24]. In the same line, we ensured that the complete homo-dimeric structure of *PfGR* was enclosed while designing the docking virtual grid. Towards this, a 3D virtual grid of 80×94×94, with x-y-z coordinates of the center of energy: 74.579, 45.32 and 92.491, respectively, was designed using Autogrid feature of AutoDock Tools, with default spacing. Highest scoring docked NTF conformations were selected depending on their strong negative free binding energies to the catalytically active pocket and the cavity harbored within the dimer interface, and monitored for possible polar connections with the amino acid residues of *PfGR*. To rule out the possibility that NTF cross-interacts with and inhibits the enzymatic activity of the host GR (*Hs*GR), similar *in silico* interaction analysis with *Hs*GR in complexation with NTF was performed. X-Ray diffraction based structural model *Hs*GR (PDB ID: 3GRS) was procured from PDB. A 3D virtual grid of 94×94×94, with x-y-z coordinates of the center of energy: 81.85, 36.101 and 18.741, respectively, was constructed, followed by interaction analysis. PyMOL Molecular Graphics System, v2.1 by Schrödinger, LLC was used for molecular visualization of the structural models of *Pf*GR and *Hs*GR, and AutoDock Vina results [29].

### 2.8 Preparation of parasite lysate

The parasite lysate was prepared in accordance with the previously described method [30]. Briefly, parasites were released from erythrocytes by treatment with 0.05% saponin, on ice for 15 min and washed with PBS. Parasite pellet obtained by centrifugation at 1300x g (5 minutes) was washed with PBS, sonicated at 4°C in PBS and stored at -20°C for further use.

### 2.9 Glutathione Reductase (GR) activity measurement

The impact of NTF on enzyme activity of *Pf*GR was measured using previously documented method **[31]**. Trophozoites parasitized erythrocytes at 8-10% parasitemia were incubated with varying concentrations of NTF for 24 h and proceeded with lysate preparation. GR activity in the parasite lysate was determined using GR assay buffer (26.5 mM K_2_HPO_4,_ 20.5 mM KH_2_PO_4,_ 1 mM EDTA, 200 mM KCl, pH 6.9) in the presence of glutathione disulfide (1 mM) and NADPH (100 μM) as substrates. The absorbance change per minute at 340 nm was recorded using a microplate reader. Specific enzyme activity of *Pf*GR was calculated in treated and control samples.

### 2.10 Estimation of Reduced (GSH) and Oxidized (GSSG) Glutathione Levels

The estimation of glutathione levels in reduced (GSH) and oxidized (GSSG) forms was performed in accordance with the previously documented protocol [32]. Briefly, trophozoites were incubated with varying concentrations of NTF for 24 h and lysate was prepared. For GSH estimation, lysate was added to assay mixture containing 0.1mM sodium phosphate buffer (pH 8) and o-phthalaldehyde (1 mg/ml in methanol) and incubated at RT for 15 min. GSH level was estimated by monitoring fluorescence change at 365_Exc_/430_Emi_ using microplate reader. GSSG was also estimated using the same method, however the lysate was incubated first with N-ethylmaleimide (NEM) in dark for 5 min, to block the GSH to GSSG oxidation. Results were plotted as redox index (GSH/GSSG) ratio.

### 2.11 Drug Combination assay

P*f*3D7 *and Pf*Kelch13^R539T^ parasites at ring stage (0.8-1%) were treated with different concentrations of NTF (2.5–40 uM) + ART (2.5-40nM) along with individual drugs, in accordance to fixed ratio method [33] for 66 h. Untreated parasites served as a control. Fractional inhibition concentrations (FICs) were computed as reported previously [34]. FIC values of <1 were interpreted as synergistic interaction.

### 2.12 Ring Survival Assay

To evaluate the efficacy of NTF on *Pf*Kelch13^R539T^ parasites, ring stage survival assay (RSA) was performed as described previously [35]. Briefly, percoll-purified schizonts were grown to early ring stage (0-3 h post-invasion), and exposed to different concentrations of NTF (6-120 µM) and DHA (700 nM) alone as well as in combination for 6 h. Following 6 h incubation cultures were washed and incubated again for 66 hr. Percent parasitemia was assessed by scoring ∼5000 RBCs in Giemsa stained smears. Concomitantly, to assess the effect of NTF with DHA on survival of DHA-resistant parasites, ring-stage (0-3h) parasites were incubated for 6 h with DHA (700 nM). Following washing, NTF (6 and 12□μM) was added and left for 66h. Percent survivability was assessed from Giemsa slides.

### 2.13 In vivo anti-malarial assay

The *in vivo* anti-malarial efficacy of NTF alone and in combination with ART was determined using Swiss albino mice infected with *P. berghei* ANKA parasites. The experiment was performed in conformity with the guidelines established by the Institutional Animal Ethics Committee (IAEC no. 35/2019.) of Jawaharlal Nehru University (JNU). Mice procured from Central Laboratory Animal Resources (CLAR), JNU were segregated randomly into four groups (n=4). Each group of mice was administered NTF (40 mg/kg in 0.5% NaCMC, p.o.), ART (60 mg/kg, i.p.), NTF (20 mg/kg) + ART (30 mg/kg) and 0.5% NaCMC (vehicle control) respectively for 5 days (Figure 6A). Parasitemia for each day was assessed from tail blood smears (Giemsa-stained). The mean survival time (in days) for mice in each group post-inoculation was calculated over a period of 45 days.

### 2.14 Statistical analysis

Statistical analysis was carried out using one-way analysis of variance (ANOVA) wherever required.

## 3. Results

### 3.1 NTF impedes intra-erythrocytic growth of P. falciparum

The *in vitro* anti-malarial activity of NTF on drug-sensitive and resistant parasites revealed that NTF impeded growth of parasites with a 50% inhibitory concentration (IC_50_) of 3.67 μM and 6.54 μM, respectively [Figure.1A (i) and (ii)]. Moreover, the effect of NTF on parasite development (66 h post-treatment) revealed that NTF treatment in *Pf*3D7 and *PfKelch*13^R539T^ parasites hampered the progression of parasites into trophozoites, instead pycnotic bodies were reported.

**Fig. 1.**
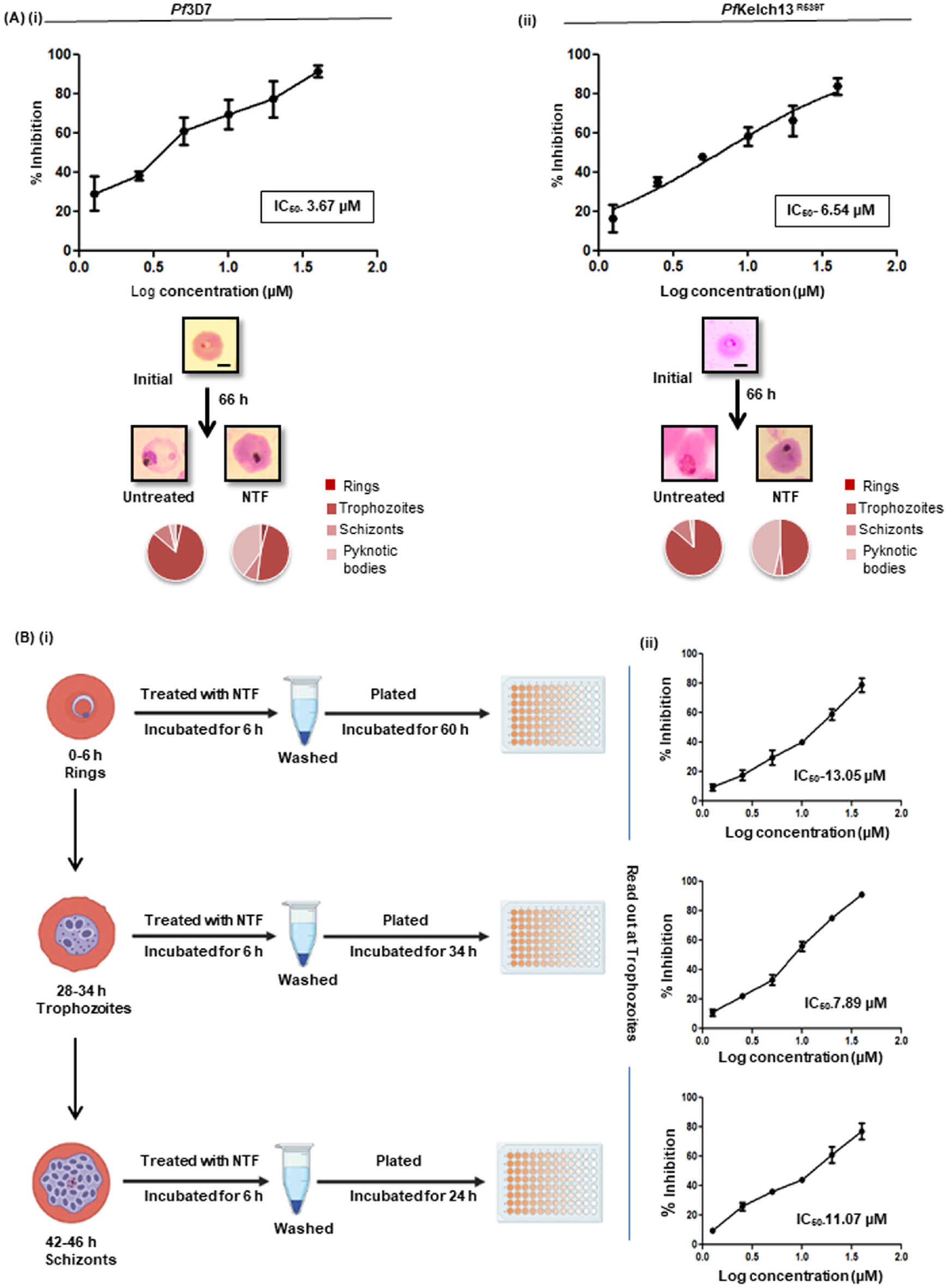
NTF inhibits the *in vitro* growth and development of ART-sensitive and resistant strains of *P. falciparum*. The concentration-dependent growth inhibitory effect of NTF on ART-sensitive *Pf*3D7 strain (i) and artemisinin**-**resistant *Pf*Kelch13^R539T^ strain (ii) as evaluated by scoring Giemsa stained parasites. Pie charts represent the proportional distribution of different intra-erythrocytic parasitic stages and pyknotic bodies at the corresponding IC_50_ values following drug exposure. (i) Schematic illustration depicts stage-specific exposure of parasite with NTF (ii) The graphs represent the percent reduction in parasitemia by NTF at each parasitic development stage, along with the respective stage-specific IC_50_ values. The result depicts the means ± SEM of three independent experiments.

### 3.2 NTF maximally targets the trophozoite stage of intra-erythrocytic cycle

The growth inhibitory potential of NTF on different intra-erythrocytic stages was checked by *in vitro* stage specific inhibition assay. A brief treatment (6 h) of the parasite with NTF at each stage depicted chemosuppresive effect of NTF in a stage-dependent manner [Figure.1B (i) and (ii)], particularly showing maximal inhibitory effect on trophozoite stage (IC_50-_7.38µM). IC_50_ values at ring and schizont stage were found to be 13.06 and 11.87 µM.

### 3.3 NTF elevates intraparasitic ROS levels

We determined the effect of NTF on liberation of free radicals and accumulation of ROS in the parasite. Untreated parasites displayed nominal ROS levels, while NTF treatment for 6 hours elevated ROS levels in both ART-sensitive and resistant parasites [Figure.2A(i) and (ii)]. As expected, ART (positive control) induced ROS in *Pf* parasites effectively. Interestingly, NTF was potent enough to generate a 1.7 fold increase in ROS levels in resistant parasites as compared to ART. Thus, NTF acts as a potent alternate inducer of ROS in both sensitive and resistant parasites.

**Fig. 2.**
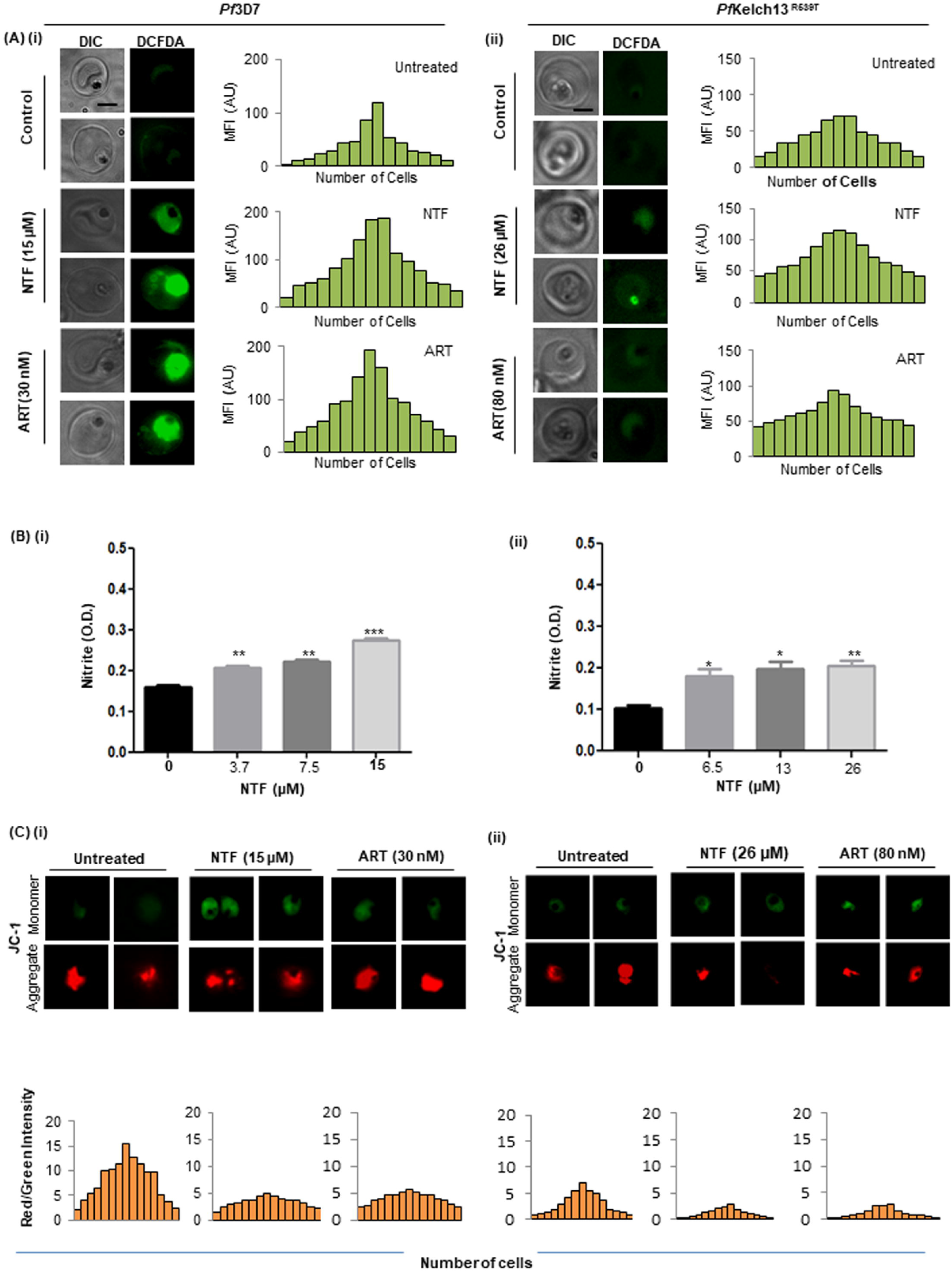
NTF induces oxidative stress and disrupts *P. falciparum* ∆Ψ_*m*_. Micrographs showing elevation in intracellular ROS levels of *Pf* 3D7 (i) and *Pf*Kelch13^R539T^ (ii) following treatment with NTF. Adjoining bar graphs depicts the mean fluorescence intensity of DCFDA for untreated and treated parasites. ART served as a positive control. **(B)** Graphs showing increased formation and release of Nitrite (NO_2_-) in the culture medium of NTF-treated *Pf*3D7 (i) and *Pf*Kelch13^R539T^ (ii) as compared to untreated control. **(C)** Fluorescence microscopic images of NTF-treated and untreated *Pf*3D7 (i) and *Pf*Kelch13^R539T^ (ii) parasites showing JC-1 aggregates (red) and monomeric JC-1 (green), indicating that NTF triggers mitochondrial depolarization. ART served as positive control. The adjoining bar graphs represent the ratio of red to green fluorescence intensity under each condition.

### 3.4 NTF elevates RNS levels in parasite

The effect of NTF on RNS production in parasites was determined by measuring the nitrite (NO^2^−) levels in the culture medium. NTF-treated parasites displayed significant increase in released nitrite, when compared to untreated control in both ART-sensitive and resistant parasites [Figure 2B (i) and (ii)). This finding suggests that NTF increases the oxidative burden in parasite by promoting RNS accumulation.

### 3.5 *NTF induces loss of* ∆Ψ_*m*_

Fluorescence microscopy of untreated parasites displayed intense red staining due to JC-1 dye accumulation inside mitochondria, indicative of their viability (Figure 2C (i) and (ii)). Interestingly, NTF-treated sensitive and resistant parasites showed diminished red staining in mitochondria and a concomitant increase in green staining in cytosol. The red:green fluorescence ratio in the untreated parasites was higher than in NTF-treated parasites, as depicted by the bar graphs. This indicates that NTF induces depolarization of mitochondria, which may contribute to programmed cell death of parasites.

### 3.6 In-silico interaction of PfGR with Nitrofurantoin

Amino acid residues of *PfGR* involved in the interaction with FAD, NADPH and GSSG, constituting the respective catalytically active sites; those involved in forming the pocket at the dimer interface; and, parasite specific insertions: “Loop 1” and “Loop 2”, are shown in Fig. 3A (i). Schematic representation of complex formation between NTF (2D structure is shown in Figure 3A(ii) and the catalytically active GSSG binding site along with the cavity formed at the intersubunit-contact of *PfGR*, are shown in Figure 3A(iv) and (v), respectively. R362 & T494 of the GSSG binding site, and D59 & E433 of the intersubunit pocket of *PfGR* were found to interact with NTF via polar interactions (H-bonds), with the acceptable bond lengths ranging from 2.8 to 3.3 Å.

**Fig. 3.**
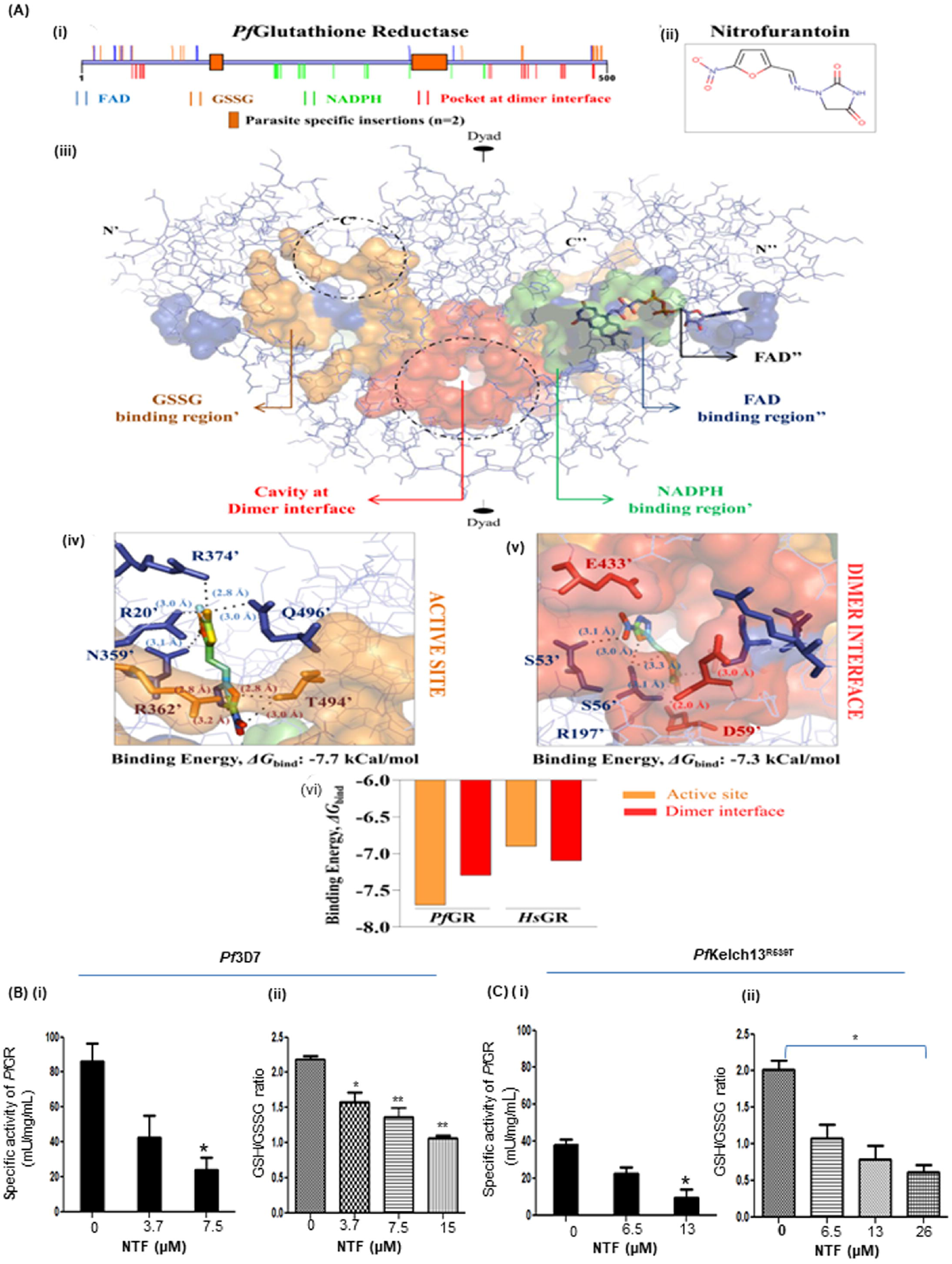
NTF targets the redox enzyme *Pf*GR. (i) **2D architecture of *Pf*GR dimer**. Amino acid residues involved in the interaction with FAD, NADPH and GSSG; and cavity at the dimer interface are shown, along with the parasite *specific* insertions: “Loop 1” (residues 126-138) in the FAD-binding domain and “Loop 2” (residues 318-350) in the protein’s central domain. (ii) **2D structure of NTF**. (iii) **3D structure of *Pf*GR dimer** highlighting the relevant catalytically active and intersubunit cavities. (iv) ***In silico* interaction analysis of NTF at the GSSG binding *active* site of *Pf*GR**. NTF interacts via H-bonds (n=8) with Δ*G*_bind_ of -7.7 kCal/mol. (v) ***In silico* interaction analysis of NTF at the intersubunit pocket of *Pf*GR**. NTF interacts via H-bonds (n=6) with Δ*G*_bind_ of -7.3 kCal/mol. (vi) **NTF selectively binds and inhibits the enzymatic activity of *Pf*GR**. *In silico* interaction analysis of *Hs*GR in complexation with NTF indicated that the compound interact with the GSSG binding catalytic site of *Hs*GR with Δ*G*_bind_ of **-**6.9 kCal/mol; and, with its intersubunit pocket with Δ*G*_bind_ of -7.1 kCal/mol. H-bonds. Bond lengths (Å) are highlighted as dashed lines. **(B)** The bar graph depicting inhibition of *Pf*GR specific enzyme activity upon NTF treatment in *Pf*3D7 parasites (*P=0.04) (ii) Graph showing GSH/GSSG ratio in NTF-treated *Pf*3D7 parasites (**P=0.004). **(C)** (i) NTF-mediated inhibition of GR specific enzyme activity in *Pf*Kelch13^R539T^ (*P=0.04). (ii) Graph showing reduction of GSH/GSSG ratio in NTF-treated *Pf*R539T parasites Data are expressed as mean values ± SEM and is representative of experiments carried out in triplicates.

NTF was found to interact via 8 H-bonds with the GSSG binding site (Δ*G*_bind_: -7.7 kCal/mol); and, via 6 H-bonds at the intersubunit pocket (Δ*G*_bind_: -7.3 kCal/mol). *In silico* interaction analysis with *Hs*GR in complexation with NTF indicated that the compound interact with the GSSG binding catalytic site of HsGR with Δ*G*_bind_ of -6.9 kCal/mol; and, with its intersubunit pocket with Δ*G*_bind_ of -7.1 kCal/mol [Figure 3A (vi)]. As a result, it was hypothesized that NTF is more likely to selectively bind and inhibit the enzymatic activity of *PfGR*.

### 3.7 NTF hampers PfGR enzymatic activity and alters GSH/GSSG ratio

Treatment of *Pf*3D7 with 7.5µM NTF significantly reduced the specific enzyme activity of *Pf*GR (∼3.67 fold) when compared to untreated control [Figure 3B(i)]. A similar dose-dependent decrease in GSH/GSSG ratio was also observed, with a 50% reduction at a concentration of 15 µM [Fig. 3B(ii)]. A similar depletion of *Pf*GR activity was also observed in *Pf*Kelch13^R539T^ strain upon treatment with 13 µM NTF [Figure 3C(i)], along with a dose-dependent decrease in GSH/GSSG ratio [Figure 3C(ii)]. Thus, our results demonstrate that NTF contributes to imbalance in redox system of parasites by inhibiting *Pf*GR, thus disrupting the cellular GSH/GSSG balance favoring enhanced oxidative stress accumulation.

### 3.8 NTF enhances the antimalarial activity of artesunate

A synergistic association of NTF and ART at few combinations with an FIC_90_ index of <1 in *Pf*3D7 strain was observed (Figure 4A). A similar synergistic interaction (FIC_50_) was observed in resistant strain, where NTF significantly potentiated the efficacy of ART (Figure 4B). Notably, in resistant strain, ART even at a dose of 10 nM was only able to reduce parasite growth upto 30%, but combining it with 5 µM NTF enhanced its inhibitory potential upto 70 %. Thus, NTF was able to potentiate the efficacy of ART in both ART-sensitive and resistant strains.

**Fig. 4.**
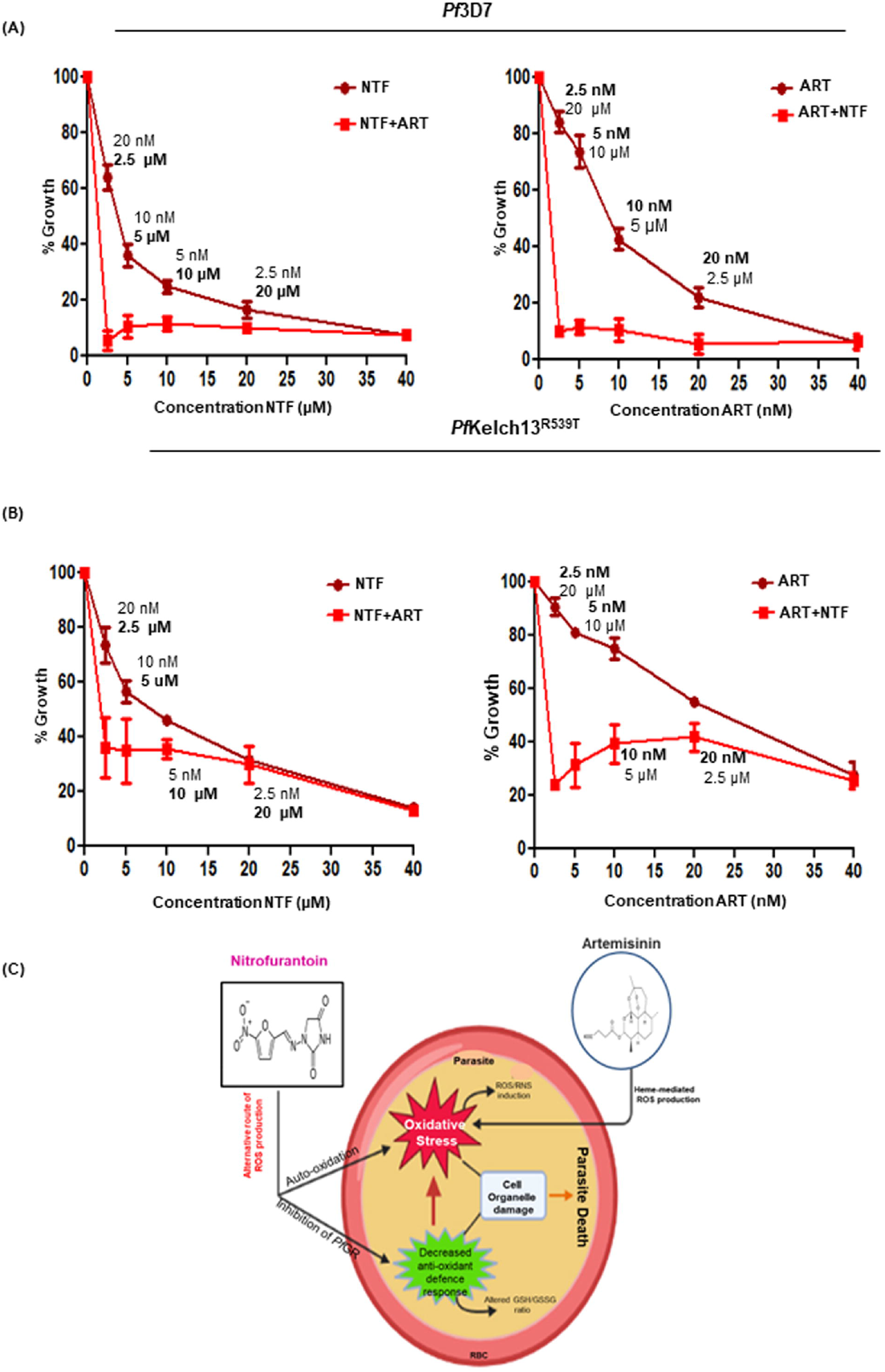
Combinatorial growth inhibitory effect of NTF and ART on *P. falciparum*. Drug combination assay demonstrating the growth inhibitory effect of NTF alone and in combination with ART on *Pf*3D7 **(A)** and *Pf*Kelch13^R539T^ **(B)**. The concentrations of each drug used in the combination are included for each data point. The graphical representation depicts growth patterns of parasites upon treatment. **(C)** Schematic representation depicting the detailed anti-parasitic mechanism of both NTF and ART.

### 3.10 NTF potentiates antimalarial potential of DHA against ART-resistant parasites

RSA on *Pf*Kelch13^R539T^ strain using DHA, NTF alone and in combination with DHA showed normal progression to healthy trophozoites in untreated parasites, while unhealthy and dead parasites were observed in NTF and DHA treated cultures [Figure 5A (i)]. After 66 h of incubation, reduced parasite survival was reported in NTF and DHA-treated resistant strain. NTF treatment resulted in >75% reduction in parasite survival [Figure 5A (ii)]. Interestingly, the combination of NTF and DHA further enhanced the efficacy of DHA by decreasing the parasitemia by 80-95% at different NTF concentrations [Figure 5A(iii)]. We also tested the inhibitory potential of NTF on resistant parasites recovered after DHA treatment. The percentage survival of DHA pretreated rings was observed in the presence of 12 µM NTF suggestive that NTF is potent in eliminating resistant parasites surviving post DHA treatment [Figure 5B (ii)].

**Fig. 5.**
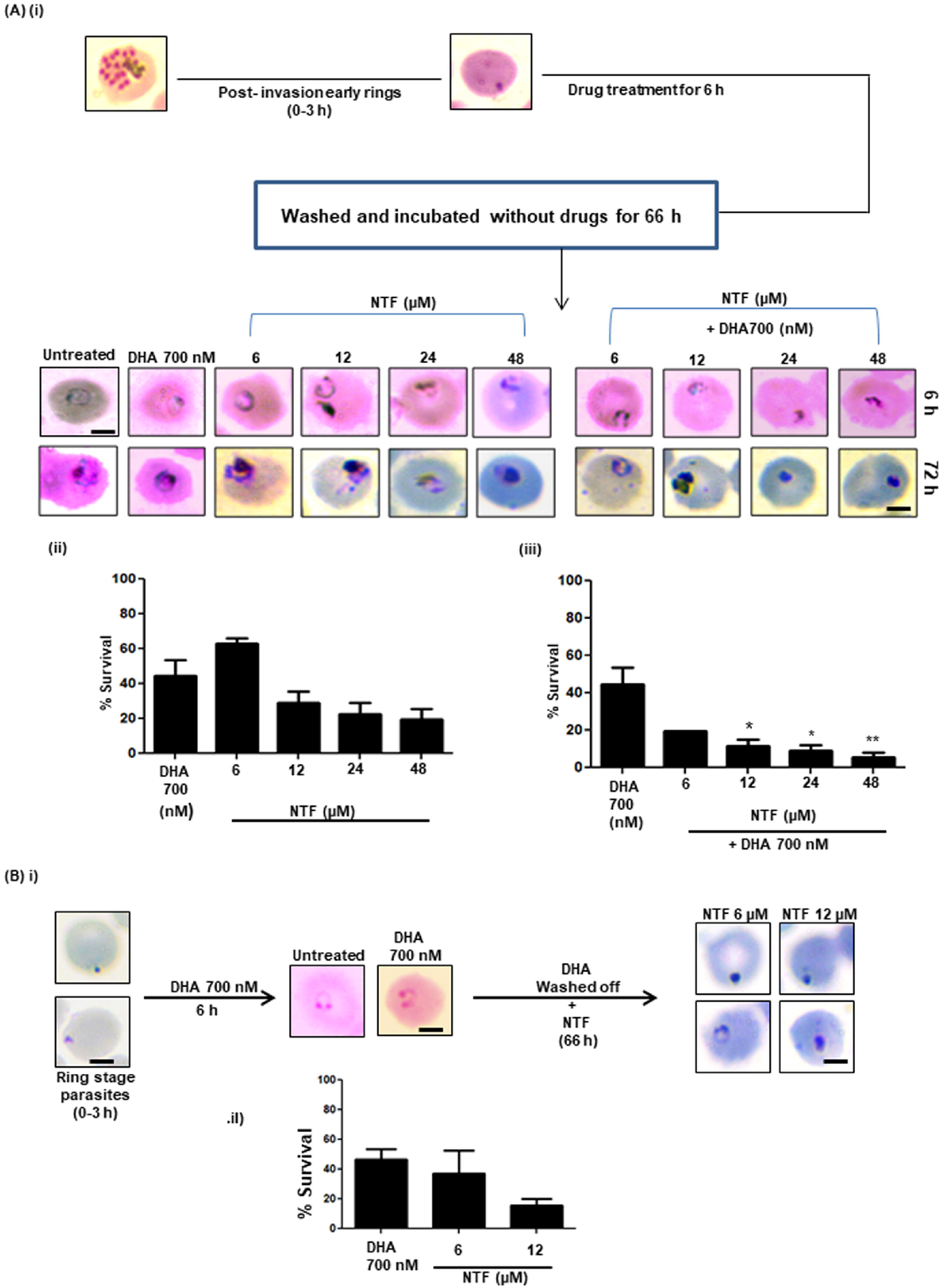
NTF potentiates the activity of DHA against ring survival of resistant parasites. **(A)** (i) Schematic representation showing light micrographs of Giemsa-stained infected RBCs incubated with DHA and NTF alone as well as in combination with DHA. Bar graphs representing percentage survival following treatment with DHA and NTF alone (ii) and in combination (iii). The combination of NTF with DHA alleviates the antiparasitic activity of DHA against ring stages of ART-resistant parasites. (*\*, P*L<L0.05, **, P<0.01) **(B)** (i) Schematic showing modified form of RSA, wherein ART-resistant ring staged parasites were treated with DHA followed by treatment with various concentrations of NTF. The surviving parasites were scored and percentage survival was calculated for each condition. (ii) NTF potently decreases the survival of DHA pretreated resistant rings when compared to DHA alone.

### 3.11 NTF exerts chemosuppressive effect in infected mice

*In vivo* efficacy of NTF in *Pb*ANKA infected mice model demonstrated a 50% reduction in parasite multiplication in NTF-treated mice as compared to vehicle control (Figure 6B & C). Moreover, NTF-treated infected mice survived for more than 13Ldays, whereas vehicle treated mice died within 6Ldays post-infection. Interestingly, combination of ART and NTF significantly decreased the parasite load as well as increased mean survival time of infected mice (Figure 6D). Mice treated with combination of NTF and ART (at half the doses) survived for more than 40 days. These preclinical results further predict NTF as a partner drug candidate to ART.

**Fig. 6.**
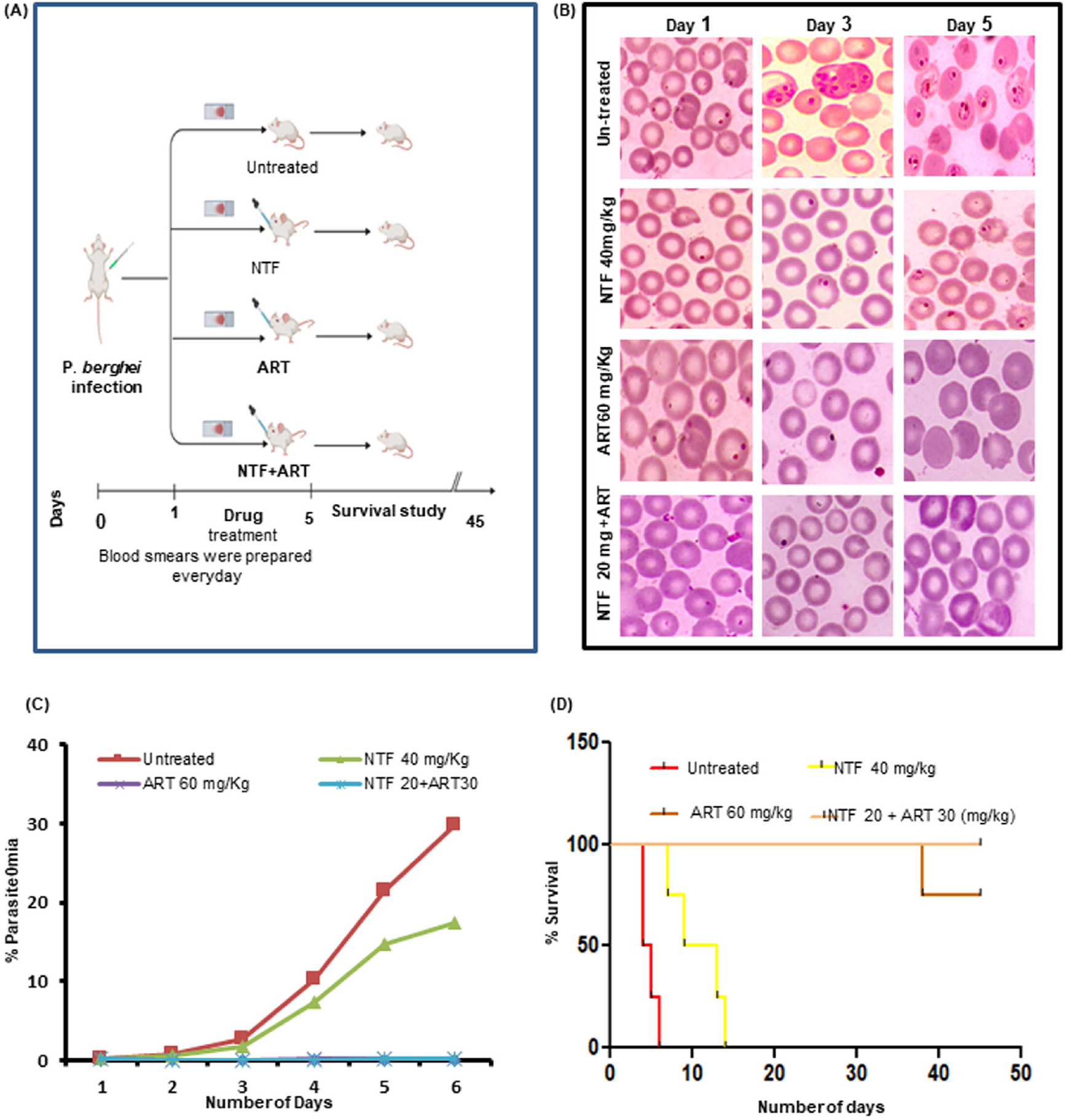
*In vivo* antimalarial potential of NTF. **(A) (i)** Schematic representation of *in vivo* drug treatment regimen. Vehicle-treated mice (n=4) served as a control. (ii) Micrographs of Giemsa-stained blood smears of control and treated mice, indicating parasite load in mice from each group. (iii) Graph showing routine percent parasitemia of untreated, NTF-treated, ART-treated and combination-treated mice. (iv) Plot showing the survival of untreated and drug-treated mice as observed for up to 45 days post-infection.

## 4. Discussion

The spread of drug-resistant *Pf* strains renders most of the currently approved antimalarials ineffective, thereby necessitating a search for new compounds. Drug repurposing has introduced multiple novel compounds that are in the process of becoming attractive antimalarial prospects [36]. Here in, we tried to reposition an antibacterial drug, NTF and investigated its anti-malarial potential against ART-sensitive (*Pf*3D7) and resistant (*Pf*Kelch13^R539T^) strains of *Pf* as well as in murine malaria *in vivo*. NTF exhibited *in vitro* potent anti-malarial effect against sensitive (3.67 µM) and -resistant (6.54 µM) *Pf* strains (Figure 1A). NTF was reported to inhibit the growth of *Pf* maximally at trophozoite stage, as evident from the stage specific inhibitory assay (Figure 1B). Interestingly, the combination of NTF and ART was found to inhibit parasite growth synergistically (Figure 4). This synergistic association between NTF with ART is being documented for the first time and it will be valuable for tackling chemoresistance, a prime hurdle in malaria eradication.

Oxidative stress-mediated parasiticidal activity is well established mechanism of traditional anti-malarials including ART. However, the evolutionary adaptability of parasite seeks to systematically counteract the anti-malarial activity mediated by ROS. The development of resistance to ART can be considered as ‘proof-of-concept’ wherein resistance has been linked to the parasite’s ability to manage oxidative stress. Reduced endocytosis of hemoglobin, leading to decreased heme mediated activation of ART, generates minimal ROS levels incapable of causing oxidative damage to parasite. Similar results were observed in our study where ROS production by ART in *Pf*3D7 was comparatively higher (2.2 fold increase w.r.t control) than in resistant parasites (1.5 fold) [Figure 2A (ii)]. Interestingly, NTF treatment resulted in enhanced accumulation of ROS in both sensitive (2.3 fold increase) and resistant (1.7 fold) parasites compared to ART [Figure 2A (i)]. Thus, our repurposed drug candidate, NTF, provides an alternate route for ROS generation in both the strains, thereby, evading the ART tolerance. Similarly, increased RNS levels were reported after NTF treatment, which further contributes to oxidative onslaught on parasites [Figure 2B (i) and (ii)]. Further, to evaluate the effect of NTF induced oxidative stress on cellular organelles, mitochondrial ∆Ψ_m_ of parasites was measured which revealed mitochondrial depolarization in NTF treated parasites [Figure 2C (i) and (ii)].

To regulate intracellular oxidative stress and redox balance *Pf* parasites utilize two redox defence systems, *i.e*., thioredoxin/thioredoxin reductase and glutathione/glutathione reductase [37], the latter one being the most prevalent [31]. The GSH/GSSG redox couple is the major cytosolic redox buffer and serves as a marker of the cellular oxidative stress and redox status. In P*f* parasites, GSH plays a vital role in the detoxification of ROS and RNS, generated mostly by hemoglobin digestion and the host’s immune system. Therefore, inhibiting GSH synthesis can serve as a potential approach in combating drug malaria. As evidenced from *in silico* study (Figure 3A), *in vitro* data also revealed inhibitory effect of NTF on the enzymatic activity of *Pf*GR [Figure 3B (i) and 3C (i)], thereby indicating its role in modulation of glutathione mediated redox balancing in *Pf*. This result supported further by significant reduction in GSH/GSSG ratio after NTF treatment in both the sensitive and resistant parasites [Figure 3B (ii) and 3C (ii)]. These results signify the efficacy of NTF in evading the oxidative stress management ability of parasites by disrupting their redox homeostasis.

The development of ART resistance linked to decreased parasites clearance has been studied using *in vitro* RSA [38]. Our RSA results demonstrated a potential inhibitory effect of NTF alone and in combination with DHA on survival of ring-stage resistant parasites [Figure 5 A]. These results indicate the efficacy of NTF against ART-resistant parasites and satisfy the prerequisites of potential drug candidate for treating drug-resistant malaria. Furthermore, we assessed the inhibitory efficacy of NTF on parasites that persisted after DHA treatment. Interestingly, NTF treatment decimated the surviving parasites (Figure 5B), thereby favoring its candidature to be a part of ART combinatorial regime.

To further validate the antimalarial activity of NTF, we assessed the *in vivo* effect in *Plasmodium* murine malaria model. NTF administration significantly reduced the parasite load and increased the mean survival time of infected mice (Figure 6). In combination-treated mice, significant chemosuppression (∼90%) was observed even at low drug dosages compared to monotherapies. In summary, NTF was found effective against both ART-sensitive and -resistant strains by exerting its anti-malarial action by accumulating oxidative stress and disrupting redox homeostasis. NTF was found to increase ROS/RNS levels as well as perturb almost all aspects of redox balance, as shown by the use of various assays such as *Pf*GR activity inhibition and GSH/GSSG ratio. Oxidative stress and redox perturbation by NTF was also observed to affect/disrupt the function of cellular function as shown by disruption in ∆Ψ_m_. Most importantly, NTF alone as well as in combination was found be active *in vivo* in murine malaria model. All these results point to future exploration of repositioning NTF as potential antimalarial compound. Moreover, our study indicates a novel approach to inhibit intra-erythrocytic development of parasite by modulating the oxidant and redox systems of *Plasmodium* and thus proposes a new way to aid in anti-malarial drug development.

## Acknowledgement

S Singh gratefully acknowledges the financial support in the form of National Bioscience Award from Department of Biotechnology (DBT; BT/HRD/NWBA/39/04/2018-19) and Indian Council of Medical Research (ICMR; NER/84/2022-ECD-I). SS acknowledges Senior Research Fellowship from ICMR (Fellowship/45/48/2019-PHA/BMS). RJ acknowledges Research Associate I fellowship from Indo-DBT (Department of Biotechnology, Government of India) and Swiss National Science foundation. AM acknowledges Council of Scientific and Industrial Research for financial aid.

---

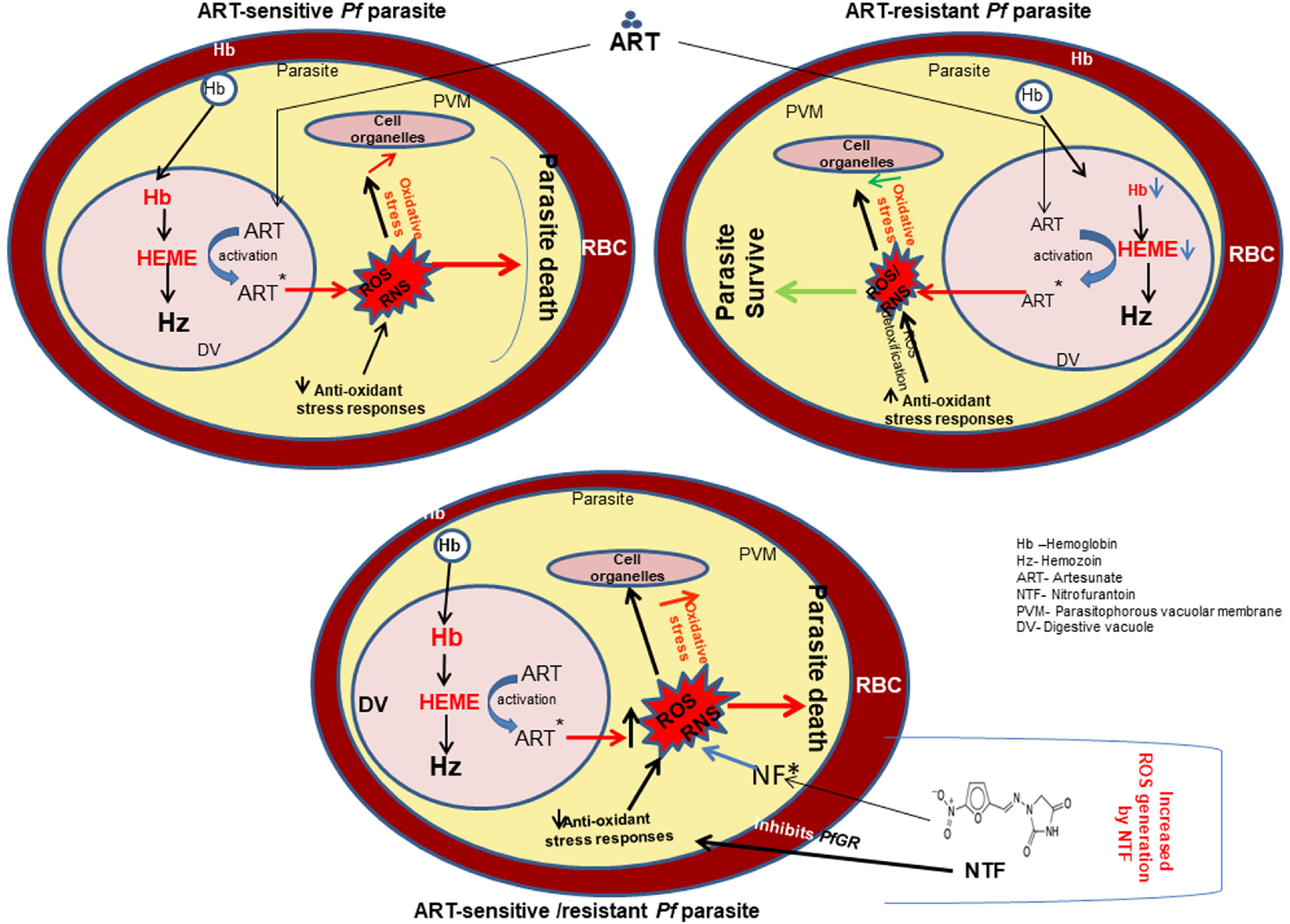

